# Conjunction or co-activation? A multi-level MVPA approach to task set representations

**DOI:** 10.1101/521385

**Authors:** James Deraeve, Eliana Vassena, William H. Alexander

## Abstract

While representing and maintaining rules in order to govern behavior is a critical function of the brain, it remains an open question as to how collections of rules - task sets - are represented in cortex. One possibility is that task sets are represented as the co-activation of representations of the simple rules from which a task set is composed. Alternatively, task sets could be encoded in a conjunctive manner as the unique combination of rules that belong to a task set. Using a novel multi-level MVPA approach in combination with fMRI, we attempted to answer both “where” and “how” task sets are represented in the brain. Subjects performed a delayed match-to-sample task using task sets composed of multiple, partially overlapping rules that governed which feature dimensions subjects should attend to, and MVPA was used to identify regions that encoded task set information. We identified voxels most relevant for classifying task sets, and, using these voxels as input to a second MVPA analysis, were able to identify regions in prefrontal cortex with activity consistent with co-active representation, while activity in visual cortex was consistent with conjunctive representation. These results highlight the utility of feature selection methods in neuroimaging analyses.

## Introduction

In cognitive experiments, participants are often required to perform tasks in which they apply simple rules, such as “*if* target is a square, *then* press left”. In everyday life, however, behavior is more complex and may be governed by collections of rules - task sets (Monsell, 2003) - that need to be selectively applied in order to achieve a goal. Consider driving your car towards an intersection: *if* you need to make a left, *then* you will need to signal this with your left indicator, AND *if* the light is red, *then* you will need to stop. Such situations and contexts are ubiquitous in daily life and require the maintenance of multiple relevant rules in working memory (WM). Moreover, adaptive behavior requires that task sets must be flexibly updated, maintained, and removed from WM in order to meet current behavioral demands.

Generally, WM and representation of rules and task sets has been reliably associated with lateral prefrontal cortex (lPFC). Early evidence for this came from studies showing impaired performance on delayed-response tasks for monkeys with lesions in this region (Goldman & Rosvold, 1970; Mishkin, 1957; Passingham, 1975). Later studies following single-unit activity in lPFC showed increased firing during retention of task-relevant information (Kojima & Goldman-Rakic, 1982; Rosenkilde, Bauer, & Fuster, 1981). Importantly, a substantial proportion of lPFC neurons was found to show selective responses to some part of the task’s events, for a large variety of tasks (Asaad, Rainer, & Miller, 2000; Funahashi & Inoue, 2000; Fuster, Bodner, & Kroger, 2000; Rainer, Asaad, & Miller, 1998). Studies using fMRI on human subjects also consistently show sustained activity during WM maintenance in lPFC (Courtney, Ungerleider, Keil, & Haxby, 1997; Clayton E. Curtis & D’Esposito, 2003; Smith & Jonides, 1999). Moreover, this delay-period activity in lPFC is sensitive to memory-influencing factors such as manipulations of delay time and the complexity and amount of memory load (Jensen & Tesche, 2002). Evidence from single-unit (Buschman, Denovellis, Diogo, Bullock, & Miller, 2012; Rigotti, Rubin, D, Wang, & Fusi, 2010) and multi-voxel pattern analysis (Cole, Etzel, Zacks, Schneider, & Braver, 2011; Dixon & Christoff, 2014) suggest that lPFC is involved principally in coding for abstract rules, possibly in a hierarchical fashion (Badre & D’Esposito, 2007).

While the above research demonstrates the involvement of lPFC in representation and maintenance of relevant task sets, the form in which they are represented, i.e., the neural code by which task-relevant information is maintained and deployed, remains an open question. It is generally accepted that, as activity propagates through the neural processing stream, the activity of individual neurons signifies increasingly abstract information. Single units in primary visual cortex, for example, respond to specific patterns of light in the visual field, while neurons located farther along the visual pathway respond to combinations of lower-level information (e.g., fusiform face area) (Amir, Harel, & Malach, 1993). The activity of neurons encoding abstract information could thus be considered a conjunctive code. In the context of rule and task set maintenance, task sets are generally considered to be more abstract, while rules govern more concrete stimulus-response contingencies (Dreisbach, Goschke, & Haider, 2007). Under the conjunctive coding hypothesis, then, task-sets should be encoded by single neurons whose activity reflects the unique combination of lower level rules. Evidence from single-unit and fMRI studies of working memory provide some evidence in support of this possibility. These studies have shown neurons coding for the identity of multiple objects/rules in lPFC, and the activation related to a particular combination is not a simple addition of the activity elicited by its parts (Nee & Brown, 2012; Sigala, Kusunoki, Nimmo-Smith, Gaffan, & Duncan, 2008; Warden & Miller, 2007). Multiple mathematical and computational models of WM have adopted an effectively conjunctive coding scheme in order to explain behavior and neural activity during WM and rule-based performance (Collins & Frank, 2013; Holroyd & Yeung, 2012; Mack, Preston, & Love, 2017; O’Reilly & Frank, 2006; Rougier, Noelle, Braver, Cohen, & O’Reilly, 2005).

Conversely, several lines of evidence suggest that representation of task sets may be consistent with *co-active* representation. Under this hypothesis, task sets are represented as the co-activation of multiple neurons, each of which codes for a single rule. Although, as noted above, neurons coding for rule combinations have been observed in lPFC, this encoding is frequently observed in parallel with neurons encoding a variety of other decision variables and combinations thereof, such as contingent motor responses (Romo, Brody, Hernández, & Lemus, 1999), stimulus-response mappings (Wallis, Anderson, & Miller, 2001), and category information (Meyers, Freedman, Kreiman, Miller, & Poggio, 2008). Computationally, recent models of WM and lPFC have suggested that lPFC may encode error representations (Alexander & Brown, 2018), with higher-level task-set information being represented in a distributed fashion as the possible errors resulting from the incorrect application of lower-level rules. Critically, in these models, rule-related error representations are shared between different task-sets when those task-sets have rules in common, i.e., representation is effectively co-active. Finally, evidence from fMRI/MVPA analysis suggest a compound representation scheme in which task sets can be decoded from the representations of simple rules (Reverberi, Görgen, & Haynes, 2012a).

The available literature thus indicates an important gap in our knowledge regarding the structure of task set representations, i.e. we lack an answer as to whether the representation of task sets in lPFC is conjunctive or co-active. Furthermore, regions in the brain beyond lPFC, such as parietal cortex and medial prefrontal cortex are also routinely implicated in the maintenance of task sets and rules (Dosenbach et al., 2007; Dumontheil, Thompson, & Duncan, 2010), yet surprisingly little research has been done to elucidate the nature of their representational schemes. Our aim in this study is thus two-fold: to identify areas encoding and maintaining task set information and to determine the nature of the representation therein. To realize these goals, we use fMRI in conjunction with a version of a delayed match-to-sample task which cues task sets with partially overlapping rules. We analyze cue-related BOLD activity during maintenance periods using a novel multi-level MVPA approach with feature selection (Deraeve & Alexander, 2018). At the first level we perform pairwise classifications of task sets and feature selection to isolate regions and indicate specific voxels that are most likely to hold discriminating task set-related information. Features selected in the first MVPA analysis are used as input features for a second analysis in which a classifier is trained to distinguish between categories composed of 1) task sets used in the first analysis and 2) novel task sets. If the representation of the task set information adheres to the conjunction hypothesis, the coding will be independent, despite the overlap of rules among task sets, and we should see a different pattern of accuracy scores on the second level than for the co-activation hypothesis, where the coding is dependent upon the separate rules. We validate this method by analyzing synthetic data sets constructed in such a way that they follow either the conjunction hypothesis or the co-activation hypothesis.

## Methods

### Participants and procedure

Data from thirty-one healthy subjects (20 female) were collected (mean age = 21.90, SD = 2.09, range 18-25). All participants were right-handed, had normal or corrected-to-normal vision and were prescreened for the absence of neurological history. We used a delayed match-to-sampling (DMTS) task having three task sets with partially overlapping rules (fig. 1). In this task, participants are given a cue lasting one second informing them of the current trial task set (color/shape CS, shape/orientation SO, or orientation/color OC). After a fixation time of seven seconds, a sample stimulus appears for one second. The subject now has to pay attention to the cued dimensions of the stimulus. After an inter-stimulus interval of three to five seconds, a target stimulus appears and subjects have to respond on how many of the cued task set dimensions the target stimulus match the sample stimulus (0, 1 or 2).

**Fig. 1.**
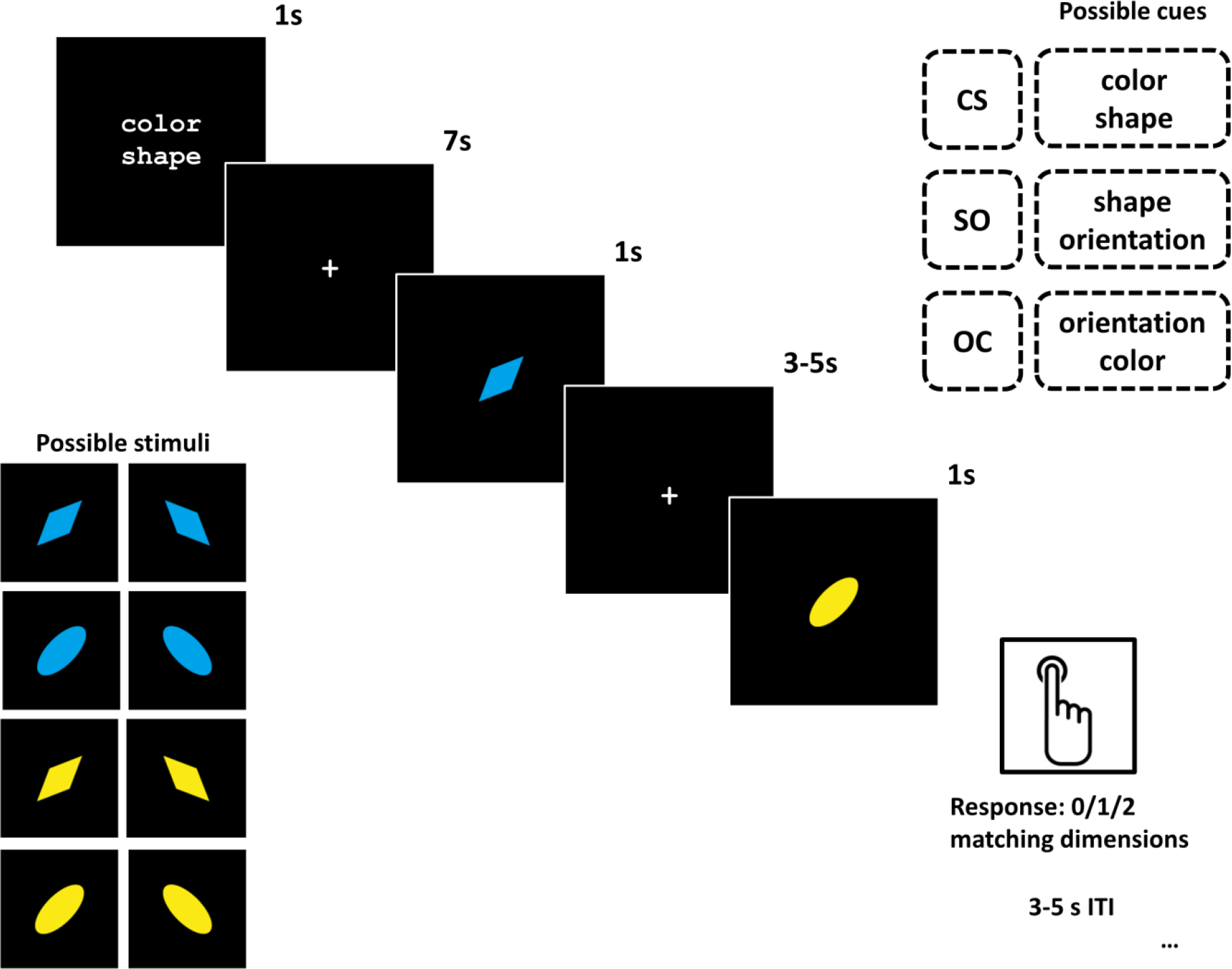
Delayed Match-to-Sample Task. At the beginning of each trial, subjects were cued with one of three possible task sets (upper right) indicating which two features of upcoming sample and target stimuli they should attend to. Subjects were required to indicating whether 0, 1, or 2 features matched. A total of 8 possible stimuli (lower right) were used in this experiment.

Our experiment also included a control condition that did not contain the rules that composed our task sets: participants are cued with either “left” or “right” and after a delay are shown an arrow pointing either left or right, and the goal was to respond whether this matches or not (fig. 3). We will refer to this task as the control task.

**Fig. 2.**
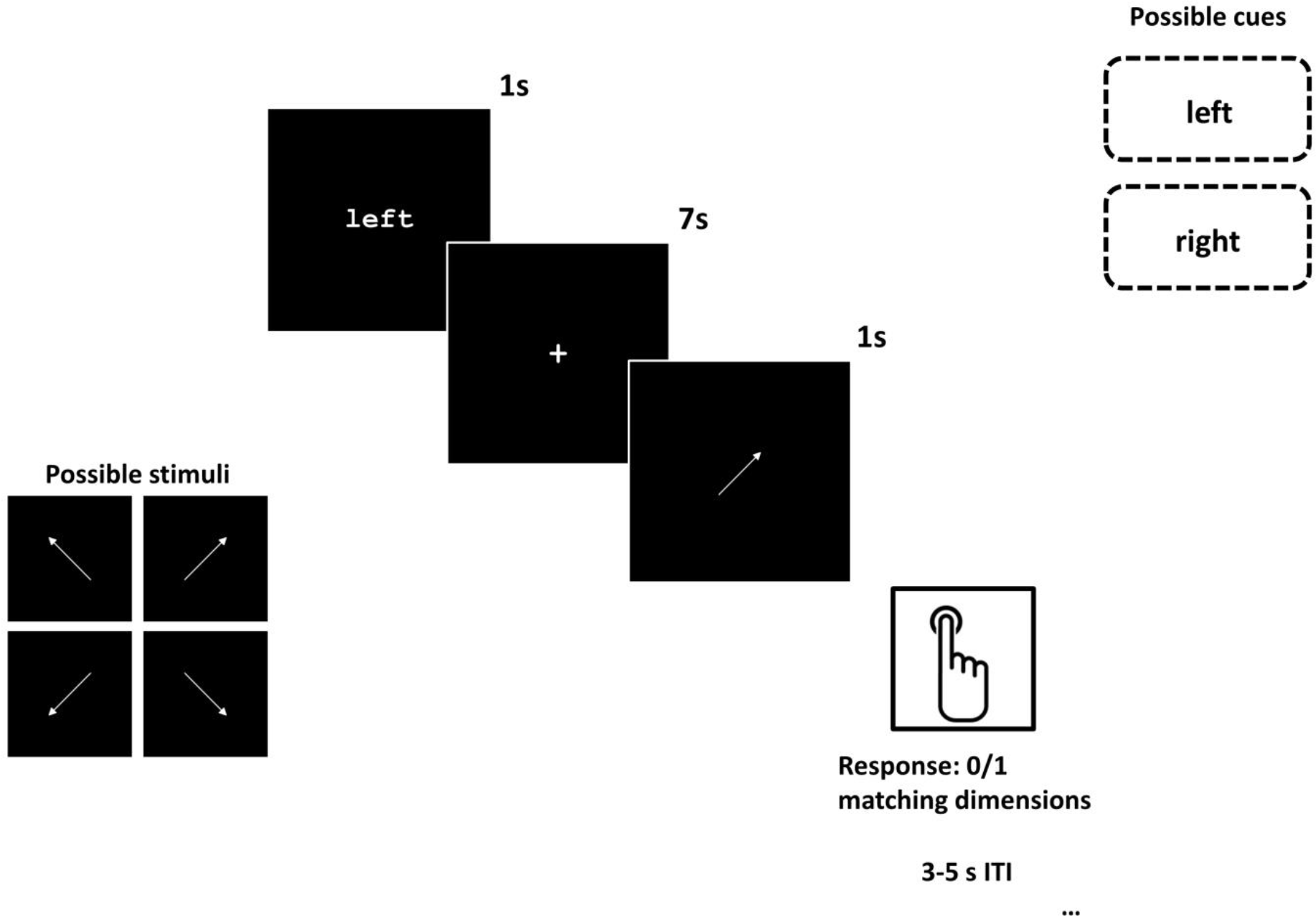
Control Task. In control trials, subjects were cued with a direction (left/right), and asked to indicate whether the target stimulus, an arrow, pointed in the cued direction.

**Fig. 3.**
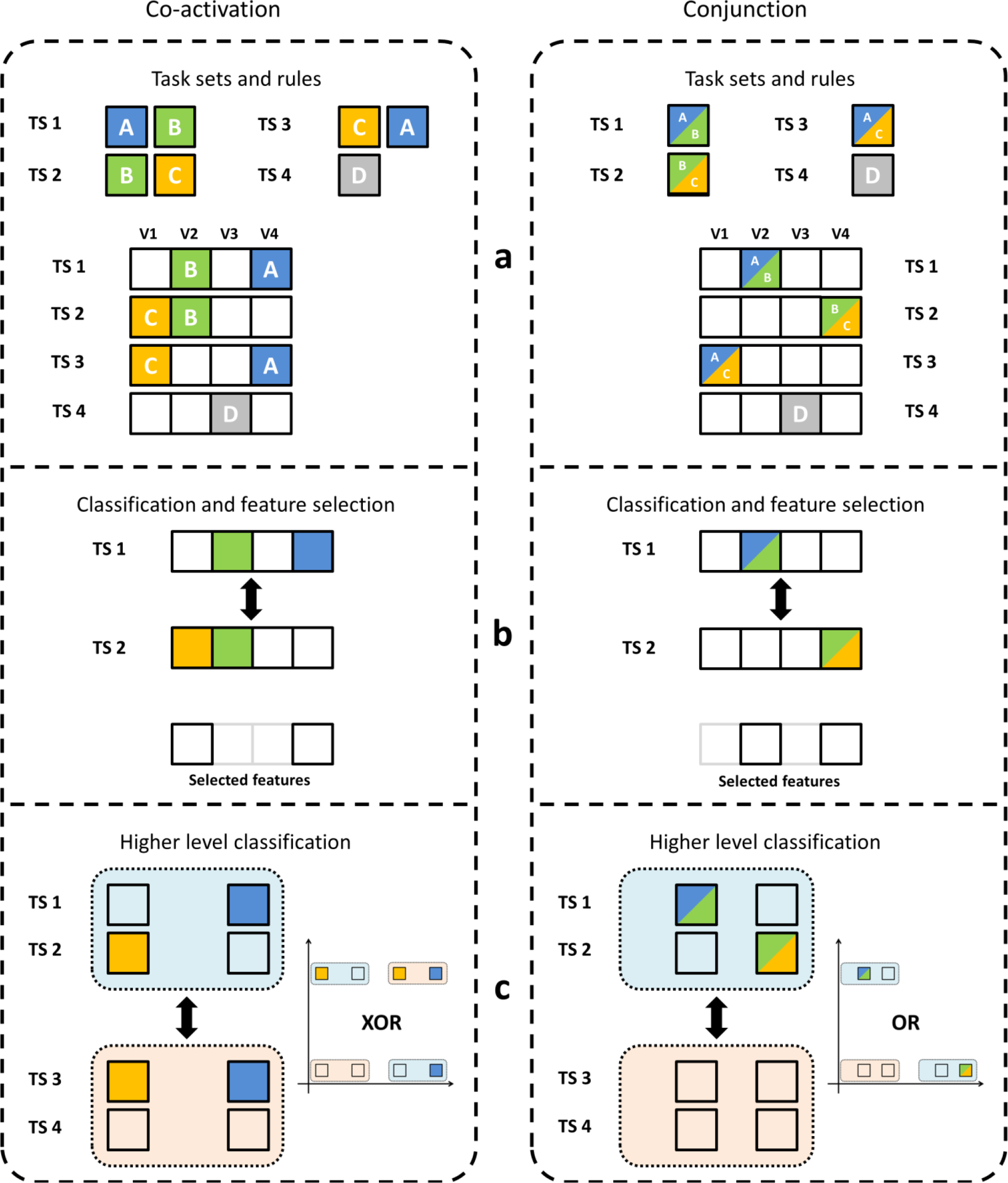
Schematic of the analysis under *co-activation* and *conjunction*. (a) Shows a simplified representation of the task sets and rules with only four features. Colored voxels carry encoding-information with their corresponding rule(s). (b) Shows how only features relevant to the classification are then selected for the next step. In (c) we take the previously classified task sets and classify them against the remaining task sets. Depending on which hypothesis is correct, this should either lead to an XOR or an OR problem. As a linear classifier can’t solve XOR problems (Minsky & Papert, 1969), successful classification provides support for the conjunction hypothesis, while classification around chance provides support for the co-activation hypothesis.

Participants were first familiarized with both tasks and then trained on a random subset of 15 (3 × 5) main task and 5 control task trials before entering the scanner to ensure they understood what was expected. The experiment itself was divided into four blocks of 37 randomized trials. Each block had 7 control task trials and 10 main task trials of each task set, resulting in a total of 148 trials across runs

### Image acquisition

Data were acquired using a 3T Magnetom Trio MRI scanner (Siemens), with a 32-channel radio-frequency head coil. The first sequence was a structural T_1_ weighted MPRAGE (176 high-resolution slices, TR = 1550 ms, TE = 2.39, slice thickness = 0.9 mm, voxel size = 0.9 × 0.9 × 0.9 mm, FoV = 220 mm, flip angle = 9°). Functional images were acquired using a T_2_ weighted EPI sequence (33 slices per volume, TR = 2000 ms, TE = 30 ms, no inter-slice gap, voxel size = 3 × 3 × 3mm, FoV = 192 mm, flip angle = 80°). Around 1280 volumes per participant were collected over an average period of 45 minutes. Image acquisition was time-locked to cue onset to easily correspond images with peak HRF.

### Preprocessing

The first 4 volumes of each functional run were discarded to allow for steady-state magnetization. Preprocessing of the data for the basic MVPA analysis and multi-level MVPA analysis was done with FSL 5.0.7, running on a Linux Ubuntu platform (version: 3.13.0). MCFLIRT (Jenkinson, Bannister, Brady, & Smith, 2002) was used for motion correction and we applied a high pass filter cutoff of 100s. PyMVPA (Hanke et al., 2009) was then used for runwise polynomial detrending and z-scoring. No additional smoothing was performed on the data. The average of the third and fourth TR after the onset of cue presentation, corresponding to peak HRF, was extracted as event of interest. Preprocessing of the data for the univariate analyses was done using SPM8 and included motion correction, co-registration of functional to structural images, segmentation and nonlinear warping to normalize the images to the MNI template image (Montreal Neurological Institute) and smoothing using a Gaussian kernel of 8 mm full width half maximum (FWHM).

### Univariate analyses

A general linear model (GLM) approach was applied to identify task set or control-specific cue-related activation. Eight task-relevant regressors were modeled. 4 regressors were used to model delay-period activity following the presentation of the task set cue. These regressors signified either Task Set or Control trials, as well as whether the eventual response was correct or incorrect. An additional 4 regressors were included at the time of the response, also corresponding with correct/incorrect task-set/control trials. The response-related regressors were modeled as event-related unit impulses, while the cue-related regressors were modeled as boxcars starting from cue onset and including the fixation period. These were then convolved with the canonical hemodynamic response function. Six additional realignment parameters were also included to account for head motion (x, y, and z translation, pitch, yaw and roll). Each block of our experiment (4 blocks total) was modeled independently, and an additional block-wise regressor was included, for a total of 60 regressors. To eliminate low-frequency noise we applied a high-pass filter (128s) and an autoregressive model was used to account for serial autocorrelation. The resulting contrast images of [task set > control] and [control > task set] were then used in a group random-effects analysis. For random effects results, the voxel-level threshold was set to 0.001 uncorrected and a whole-brain cluster-level family-wise error (FWE) correction with a p-value of 0.001 for multiple comparisons was applied.

### Basic MVPA

To further identify areas involved in working memory processes, we performed a whole-brain searchlight wherein we classified task set cue-related activation vs. control cue-related activation. This binary classification was performed using a searchlight procedure with a voxel radius of 5 as implemented in PyMVPA (Hanke et al., 2009). As classifier, we chose the default linear support vector machine (SVM) from the LIBSVM library (Chang & Lin, 2011) with 4-fold cross-validation corresponding to the scanning runs. This analysis was done on the individual subject level and afterwards resulting accuracy maps were transformed to the MNI template to allow for group-level t-tests using SPM8.

### Task set simulations & Multi-Level MVPA

In this section, we describe our multi-level MVPA analysis. First, we discuss the logic underlying the analysis stream using an example with synthetic data sets, followed by how we specifically applied the steps to the task described above.

#### Rationale

Consider three rules A, B and C. With these, we can form three combinations of task sets, each comprised of two rules A-B, B-C and A-C. Each of these task sets has one rule in common with another task set, and another rule in common with the remaining task set. Consider also a fourth rule D, which forms an independent task set which shares no rules with the other task sets. If *co-active representation* is true, task sets that share a rule will have partly overlapping activation patterns; this is not the case if *conjunctive representation* is true, due to the independent coding (fig. 3a) of task sets.

### Validation of analyses on simulated data

To ascertain whether task sets are encoded co-actively or conjunctively, we developed and validated a multi-level MVPA approach by first applying it to simple synthetic data sets. Two types of data sets were constructed that adhered to either the co-activation or conjunction hypothesis. Each data set had 300 features and 200 trials; 50 trials for each of the four task sets used in our simulated experiment. For the co-activation data sets, features 1-10 were informative of rule A, 11-20 of rule B, 21-30 of rule C and 31-40 of rule D. Features 41-300 were noise features. For task set A-B trials, informative features 1-20 (rules A and B) had values taken from a normal distribution with mean 0.8 and standard deviation 1. The same was done for task set B-C trials (informative features 21-40), task set A-C (informative features 1-10 and 21-30) and task set D (informative features 31-40). All uninformative features had values taken from a standard normal distribution (mean = 0, SD = 1). For the conjunction data sets, features 1-10 were informative for the unique conjunction of rules A-B, 11-20 of rules B-C, 21-30 of rules A-C and 31-40 of rule D. For cross-validation purposes, we constructed separate data sets in the exact same manner for both types of data. Additionally, because this is an idealized and unrealistic version of the data, we also considered data sets where informative features have mixed selectivity (i.e. they contain informative signals corresponding to other task sets/rules as well) by constructing these features from a linear combination of rule-related signals, as well as data sets with systematic noise by increasing the signal by a constant in only one of the task sets. Because the control trials have different maintenance and response requirements, we conducted an additional two simulations in which the units encoding control trials had either much higher or much lower response activity.

#### Feature selection

Performing a basic classification of task sets, as in typical MVPA approaches, yields precisely the same decoding accuracy for both types of synthetic data sets, so this in itself is not helpful in determining the representational nature. However, because of the different encoding schemes, we found that the features relevant for this classification differ contingent on the type of data set the classification was performed on (conjunction or co-activation). Consider classifying task set A-B against B-C. Under co-active encoding, the features most relevant for the classifier then correspond to the features encoding rules which do not overlap between task sets, in this case features related only to rules A and C, and not rule B, since rule B is shared between the task sets and imparts no distinguishing information. Conversely, under the conjunction hypothesis, features identified as being relevant correspond to voxels that encode the unique combination of rules for each task set, these are the features encoding A-B and B-C. Thus, extraction of the features most relevant in classifying between task sets can aid in resolving whether a given data set is encoded in a conjunctive or a co-active manner. Therefore we turned to feature selection methods as a first step in our analysis and extracted the features that were most important in classifying task set A-B against B-C for both data sets (fig. 3b).

#### Classification using selected features

While we demonstrated that feature selection results in different features depending on the data set, when dealing with a real data set where the ground truth is not known, it is still unclear whether these selected features originate from a conjunctive or co-active representation. What is known is that if the data follows a co-activation pattern, then the resulting features selected when classifying task set A-B and B-C are those features related to rules A and C, i.e. the rules that constitute the remaining task set A-C. If, however, the data follows a conjunctive pattern, then the selected features are those related to the task sets A-B and B-C themselves, and these are wholly distinct from the informative features used in the remaining task set A-C. We used this knowledge to implement a second level of analysis wherein we used the features selected from classifying A-B vs. B-C as sole input features for a second classification that uses higher-level categories composed of multiple task sets. The first higher-level category was composed out of the task sets used in the feature selection step, A-B and B-C, by giving these task set trials the same label. Similarly, the second category was created out of task sets A-C and D. When we performed a classification between these two categories using the co-activation data set, we found that the linear classifier we used was unable to distinguish between the categories. The reason for this is due to the specific input features being used, namely the previously selected features related to rules A and C. Because trials in the first category use either features related to rule A (task set A-B) or rule C (task set B-C), but not both; and trials in the second category use features related to both rules A and C (task set A-C) or none (task set D), this creates an XOR-problem which is insoluble for a linear classifier (Minsky & Papert, 1969)(fig. 3c), thus resulting in chance-level decoding accuracy. On the other hand, when we used the conjunctive data set, the features selected correspond to task sets A-B and B-C, which are only informative in the first category, but not the second, thus leading to an OR-problem which our linear classifier could easily solve, resulting in high decoding accuracy.

Interestingly, we also discovered that by performing the analysis using the least important features (those that contribute least to the classification according to the feature selection method) instead of the most important features, we discovered a pattern wherein for the co-active data set the decoding accuracy of the composite categories was higher when using the least important features than when using the most important features. We found that this was due to the presence of D rule features among the least important features. D rule features are unimportant when classifying A-B vs. B-C, but are very informative when classifying between the higher-level categories, because of task set D in one of the categories. The presence of these features changes the XOR-problem for co-activation into an OR-problem and renders it classifiable. Conversely, for the conjunctive data, using the least important features drastically reduced decoding accuracy because these are less discriminating compared to the most important features. We are therefore able to use this baseline to establish an easily distinguishable and reversed pattern corresponding to co-active representations (accuracy least important features > most important features) and conjunctive representations (accuracy most important features > least important features).

### Application of Multi-Level MVPA to fMRI

#### First level: feature selection

The first step in our multi-level MVPA analysis stream was to select voxels that discriminate well between task sets as these are assumed to be voxels that encode task set specific information (i.e. separate rules or conjunction of rules unique to the task sets being classified). We used a single-layer neural network classifier with stability selection through random subsampling and cross-validation during learning. This approach compares favorably (higher accuracy, more robust feature selection despite variance in training data, and faster execution time) to more popular feature selection approaches such as RFE-SVM and ReliefF on real and synthetic data (Deraeve & Alexander, 2018). Within the neural network, voxels serve as inputs, with output nodes for the categories. Each run lasted 30 training epochs and on every epoch weights *w*_*i*_ were batch-updated according to the delta rule:

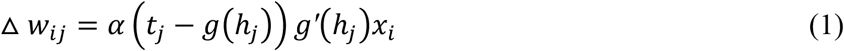

Where *α* is the learning rate (fixed at 0.01), *g*() and *g′*() are the sigmoid function and its derivative, *t*_*j*_ is the target output, *h*_*j*_ the weighted sum of the inputs, and *x*_*i*_ the input values. Weights were initialized by drawing from a normal distribution with mean 0 and standard deviation 0.01.

Data were first split into training and validation sets in a 3:1 ratio corresponding to the scanning blocks. Then the neural network fully ran for 20 replications, with random subsampling (90% of training data) on each replication to increase stability. Each epoch, performance of the neural network was assessed on the validation set to prevent overfitting. After completion, the weights from the best performing epoch on the validation set were stored, and these weights summed over all replications and cross-validation combinations. The summed weights were then sorted according to their absolute values, and the highest 5% were then selected as the most-important voxels. To establish a baseline for the end results, we also selected the 5% least-important voxels, i.e. those whose summed weights are closest to zero. This feature selection step was done for all three combinations of task sets (ts1/ts2, ts2/ts3 and ts1/ts3). An outline of the procedure can be seen in table 1.

**Table 1.**
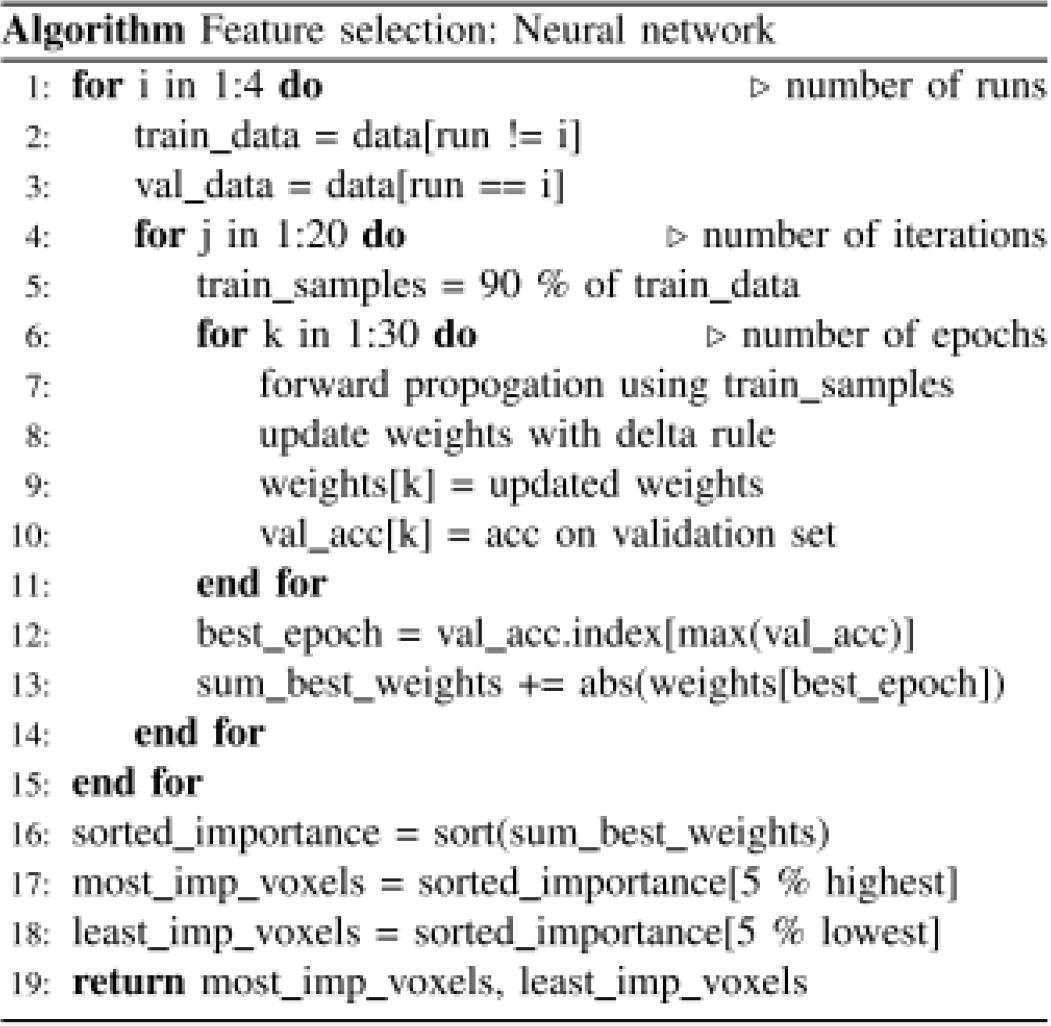
Pseudocode of the neural network for feature selection.

### Second level: classification on selected features

Two higher-level categories were created: one consisting of the task set trials of the feature selection step used to identify the features used here as input; the other from the remaining task set trials which had not yet been used and the control task trials. The data was divided in combinations of training and test sets, in a 3:1 ratio as was the case for the feature selection step. Furthermore, because of an imbalance between the number of task set trials and control trials, we applied resampling with replacement for all task set and control trials, with the number of samples equal to the mean of the number of task set trials. This resampling procedure was replicated 10 times and on each replication a linear SVM was used to classify between the two categories, after which performance results were averaged across replications and cross-validation combinations.

### Searchlight implementation

We implemented our multi-level MVPA approach in a whole-brain searchlight procedure to examine how various parts of the brain encode these task sets. All steps were performed in the native subject space and resulting accuracy maps were later normalized and assessed at the group level. Since the analysis consists of multiple levels, we found it necessary to reduce computational cost by skipping every other voxel in the searchlight, leading to a geometrical reduction in the total number of spheres created. The voxels that were left out were assigned a value equal to the mean accuracy of all the searchlight spheres they belonged to. This had the added benefit of smoothing out the final results. A radius of 5 voxels for the spheres (approximately 500 voxels) was chosen to ensure adequate remaining voxels after the initial feature selection step.

## Results

### Behavioral results

To confirm that participants were able to perform the task, and to exclude subjects from further analyses who were unable to perform the task, we tested subject response accuracy. Due to the relatively low number of trials per condition in our task (40 trials per task set condition), we adopted an exclusion criteria of no more than 20% errors in any condition in order to ensure a sufficient number of correct trials for subsequent analyses. After applying this exclusion criterion, seven subjects were removed from the initial pool. Over all task set trials, remaining subjects had a mean accuracy of 90.94% (SE 1.43%), and an accuracy of 92.70% (SE 1.26%) for control trials, indicating that, as a group, subjects were able to perform the task at levels comparable to those reported in studies using similar compositions of task rules (Cole et al., 2011; Reverberi, Görgen, & Haynes, 2012b).

### Univariate results

After preprocessing, we excluded one additional participant from further analyses due to excessive motion (multiple spikes of > 2mm). In order to provide initial evidence that regions of the brain implicated in working memory tasks were recruited by our task, we carried out a mass-univariate analysis to identify working memory regions and regions important in representing and maintaining task sets. Working memory tasks have been associated with increased BOLD activity during maintenance periods in a distributed network of regions, including lateral PFC, anterior cingulate cortex (ACC), and parietal cortex (Dumontheil et al., 2010). Our univariate analyses therefore investigated differential BOLD activity related to trials with low working memory demands (control trials) with high WM demands (task set trials) during delay after cue presentation (figure 5). Coordinates reported here and in the following sections always use the MNI coordinate system (x, y, z; mm).

For the [task set > control] contrast, we found significant activation (voxel threshold p<0.001, FWE-corrected at cluster level p(FWE)<0.001) in bilateral occipital cortex (OCC) (p(FWE)<0.001, −6, −60, 0, voxel extent = 16897) left lateral prefrontal cortex (lPFC) (p(FWE)<0.001, −52, 32, 10, voxel extent = 496), left postcentral gyrus (p(FWE)<0.001, −50, −10, 20, voxel extent = 919) and posterior cingulate gyrus (p(FWE)<0.001, 20, 30, 32, voxel extent = 968). The reverse contrast of [control > task set] resulted in significant (voxel threshold p<0.001, FWE-corrected at cluster level p(FWE)<0.001) activation of bilateral inferior parietal lobule (left: p(FWE)<0.001, −38, −42, 48, voxel extent = 2269; right: p(FWE)<0.001, 56, −54, 48, voxel extent = 385). Our univariate results thus reveal several areas traditionally involved in WM such as parietal cortex and lPFC (figure 4).

**Fig. 4.**
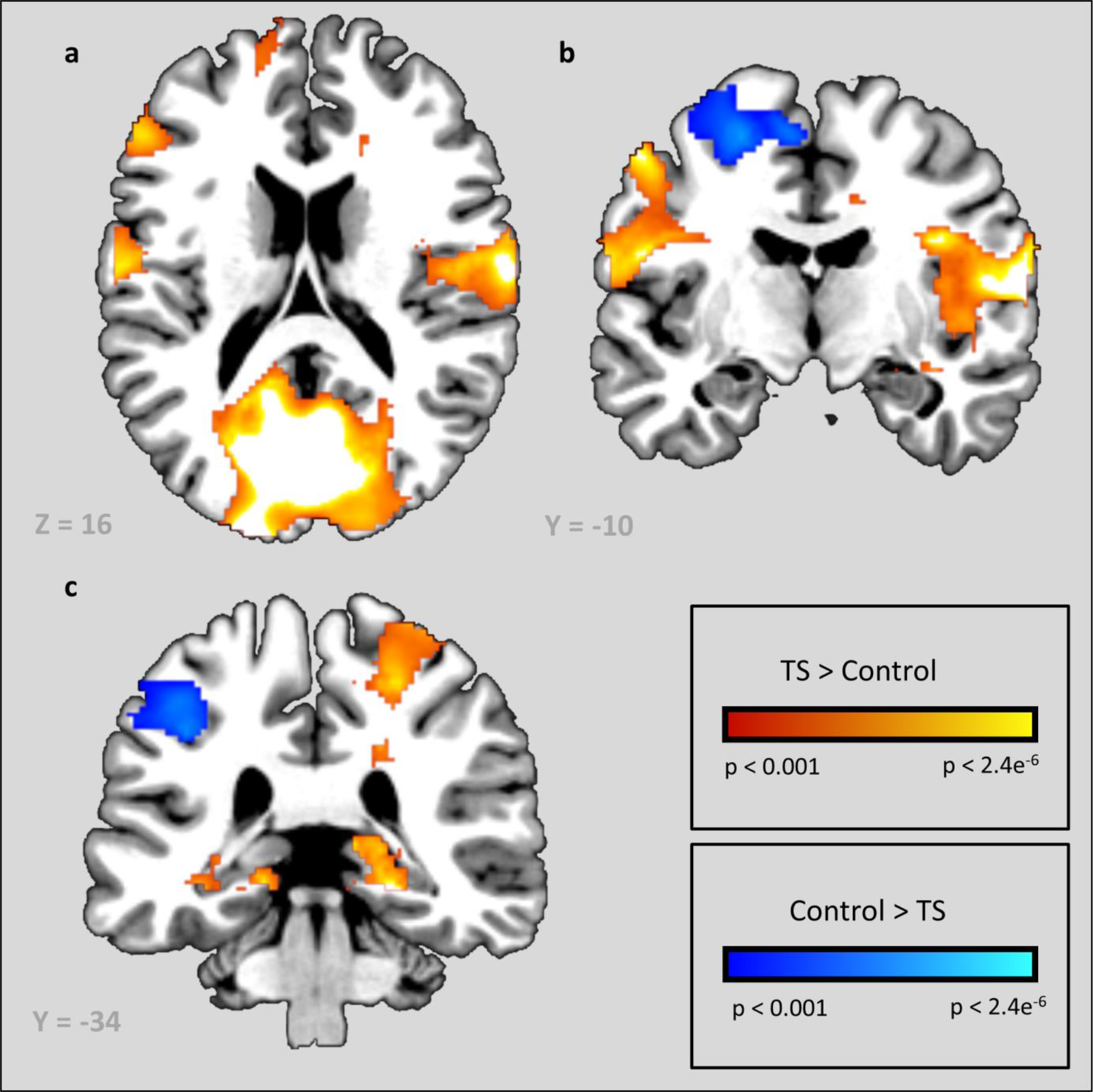
Univariate results for two contrasts. Activation clusters for [task set > control] are located in bilateral occipital cortex (OCC), left lateral prefrontal cortex (lPFC) and the left postcentral gyrus. Significant activation for [control > task set] was found in bilateral parietal cortex extending dorsally.

### Basic searchlight results

In order to provide additional evidence that regions associated with WM were involved in our task, we performed a whole-brain searchlight multi-voxel pattern analyses (MVPA) in which an SVM classifier was trained on the categorization of task set trials vs control task trials. This analysis was performed at the subject level after following the MVPA preprocessing steps as described in the methods section. After the searchlight, the resulting accuracy maps were reduced by a constant value equal to the chance level for the classification and put into a common template space. A t-test at the group level was then performed using SPM8 to identify areas exhibiting above-chance classification accuracy. Due to high statistical test values, we observed substantial trans-regional cluster formation at the typical threshold of p < 0.001. In order to ensure the specificity of anatomical regions, we therefore used a more stringent threshold of p < 0.0001, whole-brain FWE-corrected at the voxel level to identify anatomically-distinct clusters of activity. This test revealed significant (threshold p < 0.0001, FWE-corrected at voxel level) clusters similar to those found in the univariate contrasts including bilateral occipital cortex (left: p(FWE)<0.001, −18, −96, 12, voxel extent = 1606; right: p(FWE)<0.001, 14, −98, 4, voxel extent = 936), in bilateral precentral cortex (left: p(FWE)<0.001, −38, −4, 44, voxel extent = 10125; right: p(FWE)<0.001, 38, −4, 48, voxel extent = 486), and precuneus (p(FWE)<0.001, 18, −62, 46, voxel extent = 589). Additionally, we found left precentral cortex extending into lPFC (BA 6 and 9) (figure 5).

**Fig. 5.**
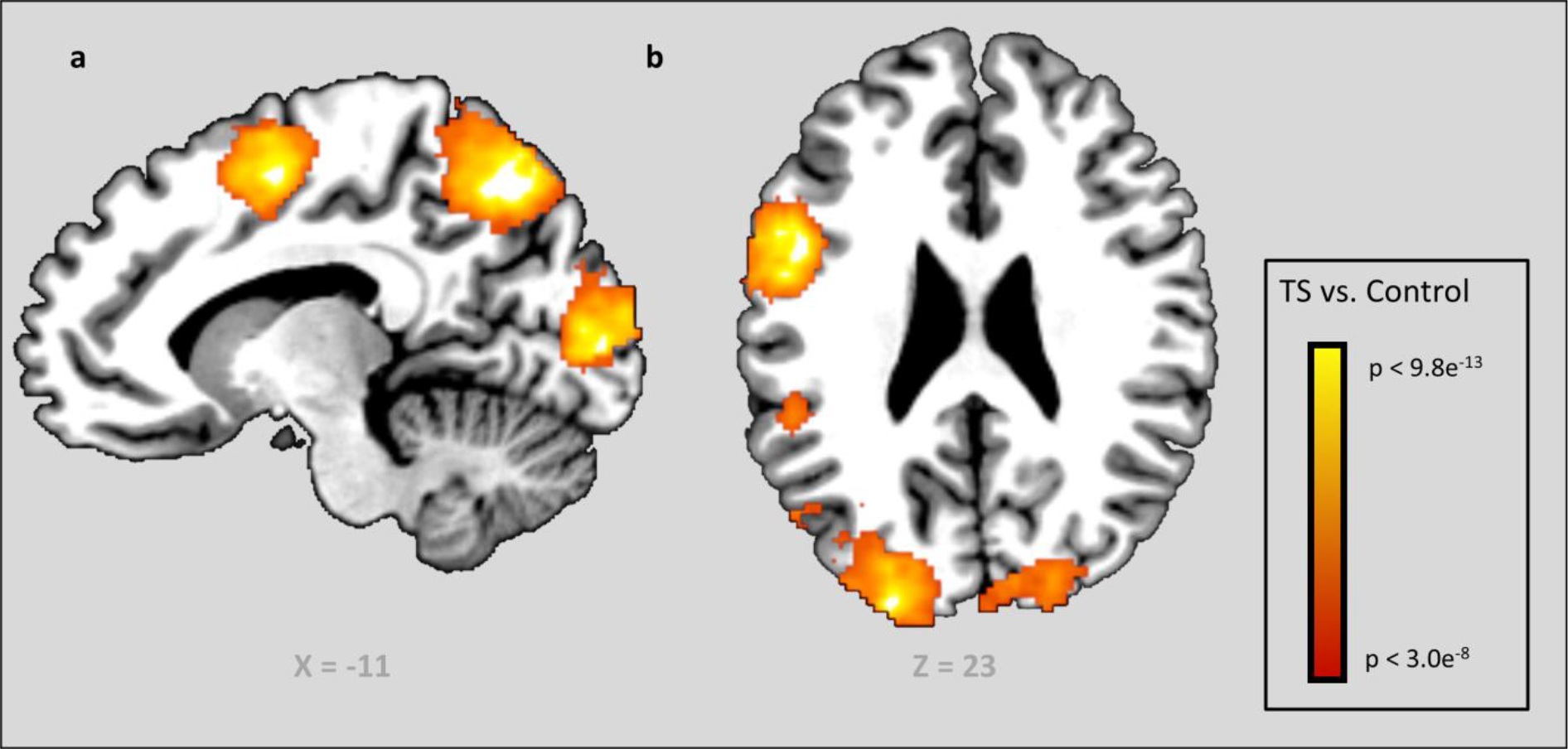
Searchlight results for classifying between task set and control trials. Clusters with significant (p < 0.0001, FWE-corrected) above-chance classification accuracy for [task set vs. control] were found in bilateral OCC, bilateral precentral cortex extending into lPFC and precuneus.

The results of this searchlight analysis suggests that, during maintenance periods, a broad array of regions represent task-set information, including those that have previously been implicated in WM such as lateral PFC, and parietal cortex. Together with our univariate analyses, these results provide multiple lines of evidence that our 2-feature DMTS task elicited neural correlates of WM. We therefore turn to our specific analyses related to the nature of information representation during maintenance.

### Simulation results

To establish that the type of representation scheme – conjunction or co-activation – can lead to differentiable patterns of classification accuracy, we simulated the multi-level MVPA analysis described in Methods on synthetic data sets which were generated according to either the conjunction or the co-activation account. Simulations were conducted for 100 independently generated data sets for each representation scheme. At the first level of the analysis, we performed a pairwise classification of two task sets, after which relevant features were extracted using the learned weights of the neural network classifier. This pairwise classification resulted in a high pairwise decoding accuracy (mean = 92.8%, SE = 1.64%) and the feature selection method was able to consistently extract all the important voxels for both conjunction and co-activation representation data sets. Note that accuracy results at this level were exactly the same for both conjunction and co-activation data sets, but differed on the sets of selected features due to shared rule-related activation among task sets for co-activation but not for conjunction. The features selected during this level were then used as input for the second level analysis.

At the second level, results show a dissociable pattern of performance based on whether the data set was generated under a conjunction or co-activation representation scheme (figure 7). If co-activation is true, using the most-important voxels of the pairwise classification as input features leads to classification performance around chance (mean = 50.21%, SE = 0.18%), and the classification using the least-important voxels as input features leads to higher accuracy (mean = 59.66%, SE = 0.83%). If the data sets were generated according to the conjunction account, then using the most-important voxels as input features at the second level, yields much higher accuracy than using the least-important voxels as input (respectively mean = 77.3%, SE = 0.22% vs. mean = 55,56%, SE = 0.52%) (see figure 6). This qualitatively different pattern shows that difference scores of accuracies of [most - least] provide a means of determining whether areas in the brain represent task sets in a conjunctive or co-active manner: under the co-activation account, classification accuracy at the second level for the most-relevant features should be lower than accuracy for the least relevant features, while the reverse is suggested for the conjunction account. Note that, using the least-important voxels, classification accuracy for both co-active and conjunctive representation is above the 50% chance level. The reason for this is that control task-related voxels are more likely to be among the least-important voxels, thus making the problem linearly separable for the least-important voxels baseline.

**Fig. 6.**
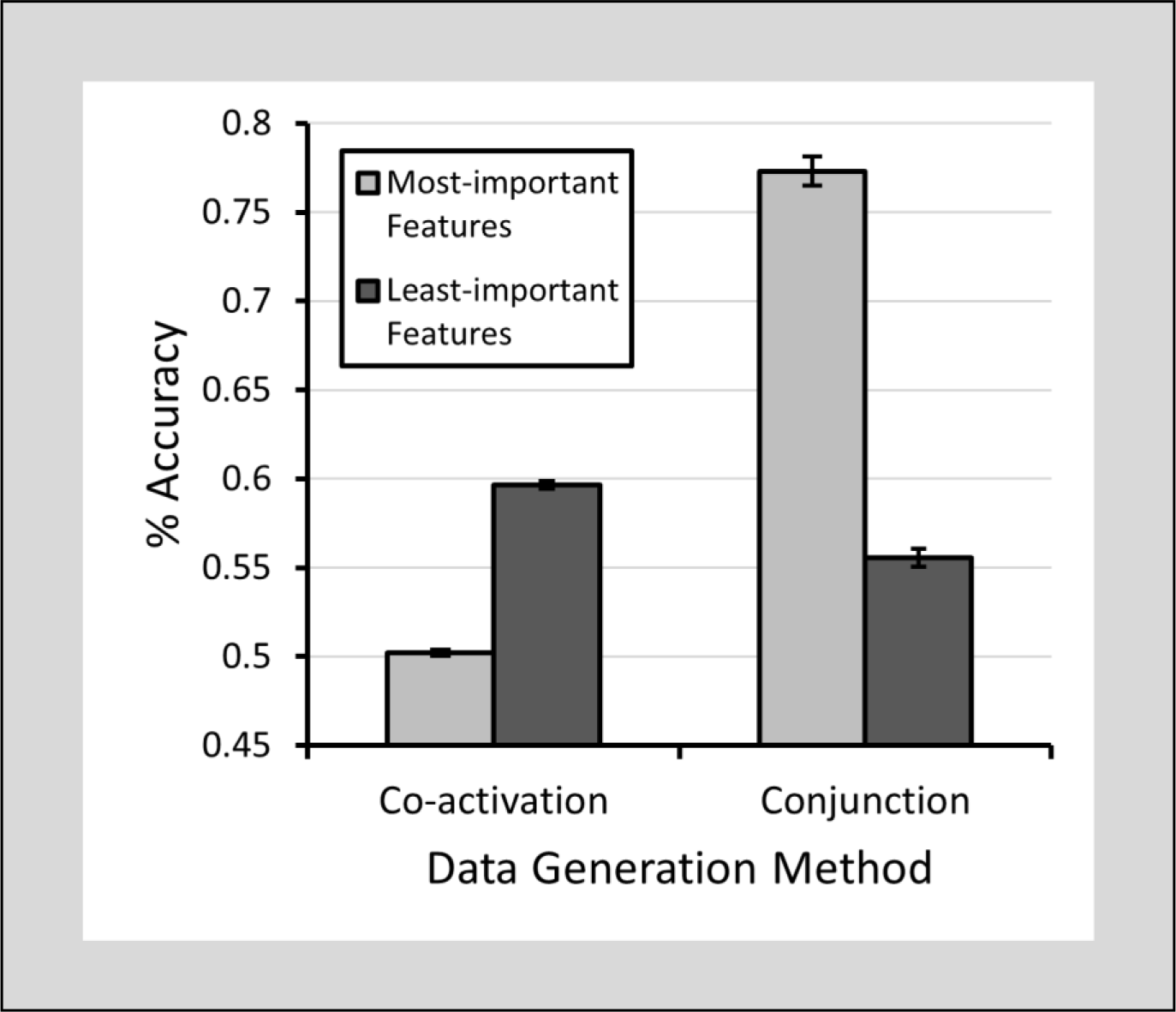
Results of the multi-level MVPA analysis on synthetic data sets. Performance of the classifier at the second level of analysis if co-activation or conjunction is true. Error bars represent standard errors of the mean (SE).

Additionally, because of the ideal nature of the data sets we constructed, we decided to investigate how robust the effect was under the “messy” conditions of systematic noise and mixed selectivity. For the systematic noise condition, we created the same data sets as above, but we added a constant noise component to all features for one set of task set trials by adding a mean shift of 0.2. Running the multi-level MVPA-analysis, we found a similar pattern of performance as above, showing it to be robust to systematic noise. A differential effect of systematic noise on the pairwise classification scores was found, where the co-activation data sets yielded higher accuracies than the conjunction data sets (respectively mean = 96.94%, SE = 0.13% vs. 90.31%, SE = 0.19%). At the second level, we again found the pattern where decoding accuracy for the co-activation data sets is higher when using the least important features compared to the most important (respectively mean = 60.83%, SE = 0.61% vs. 56.27%, SE = 0.33%), and this is reversed for the conjunction data sets where the most important features yield higher decoding accuracy between the higher level categories (mean = 75.75%, SE = 0.26% vs. 56.97%, SE = 0.25%) (figure 8a).

For the mixed selectivity condition, the informative features were modeled as a linear combination of rule-related signals to simulate sensitivity of voxels to multiple rules (co-activation) or rule combinations (conjunction). Whereas in our original data sets, each informative feature was only active during implementation of one specific rule or rule combination, in this version, informative features can respond to multiple rules or rule combinations. Specifically, for a task set A-B trial in the co-activation data set, rule A features would have values taken from a normal distribution with a mean of 0.8 (rule A sensitivity) + 0.2 (minor rule B sensitivity) and standard deviation of 1. A rule C feature in the same condition would have a value taken from the same distribution but with mean 0.2 (minor rule A sensitivity) + 0.2 (minor rule B sensitivity). This was done similarly for conjunction data sets, where features were sensitive to a mixture of rule combinations. In this manner, features still retained increased sensitivity to one of the rules or rule combinations but are also allowed to exhibit activation due to other relevant rules or combinations. Pairwise classification scores (mean = 83.95%, SE = 0.22%) were high and exactly the same for both conjunctive and co-activation data sets. At the second level, we found the same pattern as before, whereby the co-active data sets using the least important features yielded higher accuracy compared to using the most important features (respectively mean = 57.39%, SE = 0.73% vs. mean = 55.50%, SE = 0.22%). This pattern was again reversed for the conjunctive data set which showed increased performance for the most important features compared to the least important (respectively mean = 70.61%, SE = 0.24% vs. mean = 53.84%, SE = 0.43%) (figure 7b). This suggests that voxels coding for multiple rules or combinations of rules does not pose a problem to the multi-level MVPA analysis used here.

**Fig. 7.**
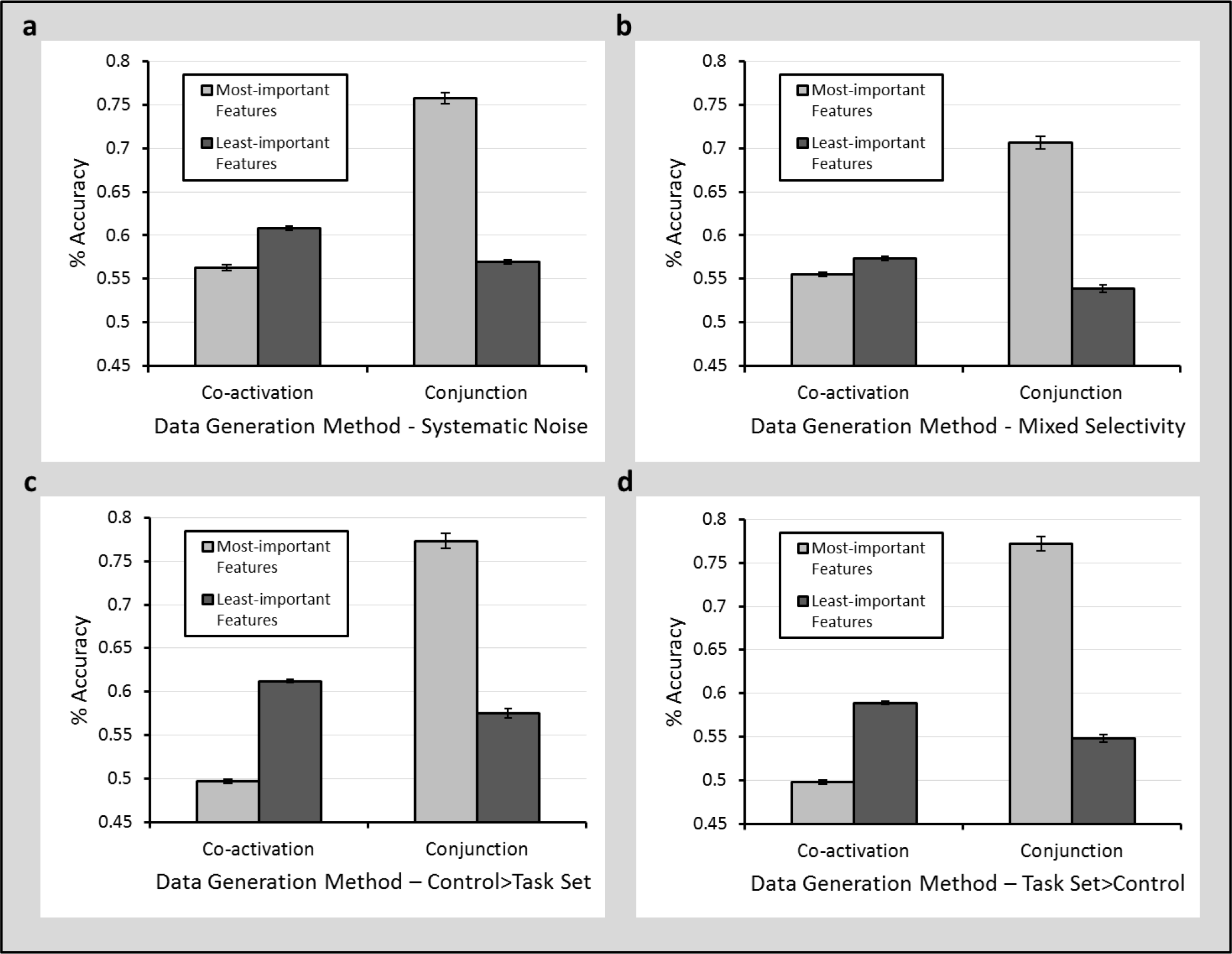
Results of the multi-level MVPA analysis on synthetic data sets under “messy” conditions. Performance of the classifier at the second level of analysis if co-activation or conjunction is true under conditions of a) systematic noise, b) mixed selectivity, c) higher control trial activity, and d) higher task set activity. Error bars represent standard errors of the mean (SE).

Finally, as indicated by our basic searchlight results in which classification accuracy was significantly above chance over large regions of cortex, the strength of the BOLD signal on control trials was likely quite different relative to task set trials. To verify that changes in the strength of the BOLD signal in control trials do not impact our analyses, we conducted two additional simulations in which the mean activity of task set trials was 1) lower (mean task set = 0.8, mean control = 1.0) or 2) higher (mean task set = 1.0, mean control = 0.8) than control trials. Results of these simulations demonstrate that, regardless of the relative signal strength of task set vs. control trials, the rationale underlying our analyses still holds. For the simulation in which control trial activity > task set activity, second-level classification accuracy for the least important features was higher than for the most important features (respectively mean = 61.21%, SE = 0.84% vs. mean = 49.70%, SE = 0.22%) under co-active coding, while the reverse is true under conjunctive coding (respectively mean = 57.53%, SE = 0.53% vs. mean = 77.33%, SE = 0.18%) (figure 7c). If task set activity > control activity, then under co-active coding the same result occurs in which the least important features result in higher accuracy at the second level of analysis than the most important features (respectively mean = 58.89%, SE = 0.83% vs. mean = 49.75%, SE = 0.24%), and under conjunctive coding we again get the reverse (respectively mean = 54.79%, SE = 0.42% vs. mean = 77.19%, SE = 0.18%) (figure 7d).

### Multi-level MVPA searchlight results

In order to identify brain regions with activity consistent with either co-activation or conjunction we conducted a full-brain searchlight analysis using our multi-level MVPA analysis (Fig. 3). Since our earlier analyses already identified key regions relevant to task set representations, converging results here would not be surprising. However, these analyses were agnostic regarding the nature of the representation. The added value of the multi-level MVPA analysis then lies in its ability to determine whether classification accuracy in these regions is consistent with either the co-activation or conjunction of task set rule representations.

Based on our simulation results, in order to identify a region as engaging in either conjunctive or co-active task-set representation, the region must meet the following criteria: first, it should be possible to decode pairwise task-set information from that region, and second, classification accuracy using voxels identified as most relevant for pairwise classification should be greater than classification accuracy for the least relevant voxels (conjunction) or vice-versa (co-activation). Accordingly, in order to identify regions of the brain involved in encoding task-sets, following the first level of our multi-level MVPA analysis for the 3 pairwise task-set categorization problems (OC vs SO, OC vs CS, SO vs OC), an accuracy map was created for each subject using the median accuracy of all 3 pairwise classification analyses (figure 8). Subject accuracy maps were converted to a common template space, and a group t-test was conducted to identify regions that encoded task-set information. In agreement with a substantial body of WM literature, above-chance classification for task sets was observed in left ventrolateral PFC (peak voxel −46, 20, 20, p<0.001) and parietal cortex (peak voxel −24, −48, 68, p=0.011), bilateral visual cortex (peak voxel 22, −98, 8, p<0.001), and medial PFC(peak voxel 12, 56, 12, p<0.001). The results of this analysis largely recapitulate our previous analyses showing regions that successfully encode task-set vs control trials.

**Fig. 8.**
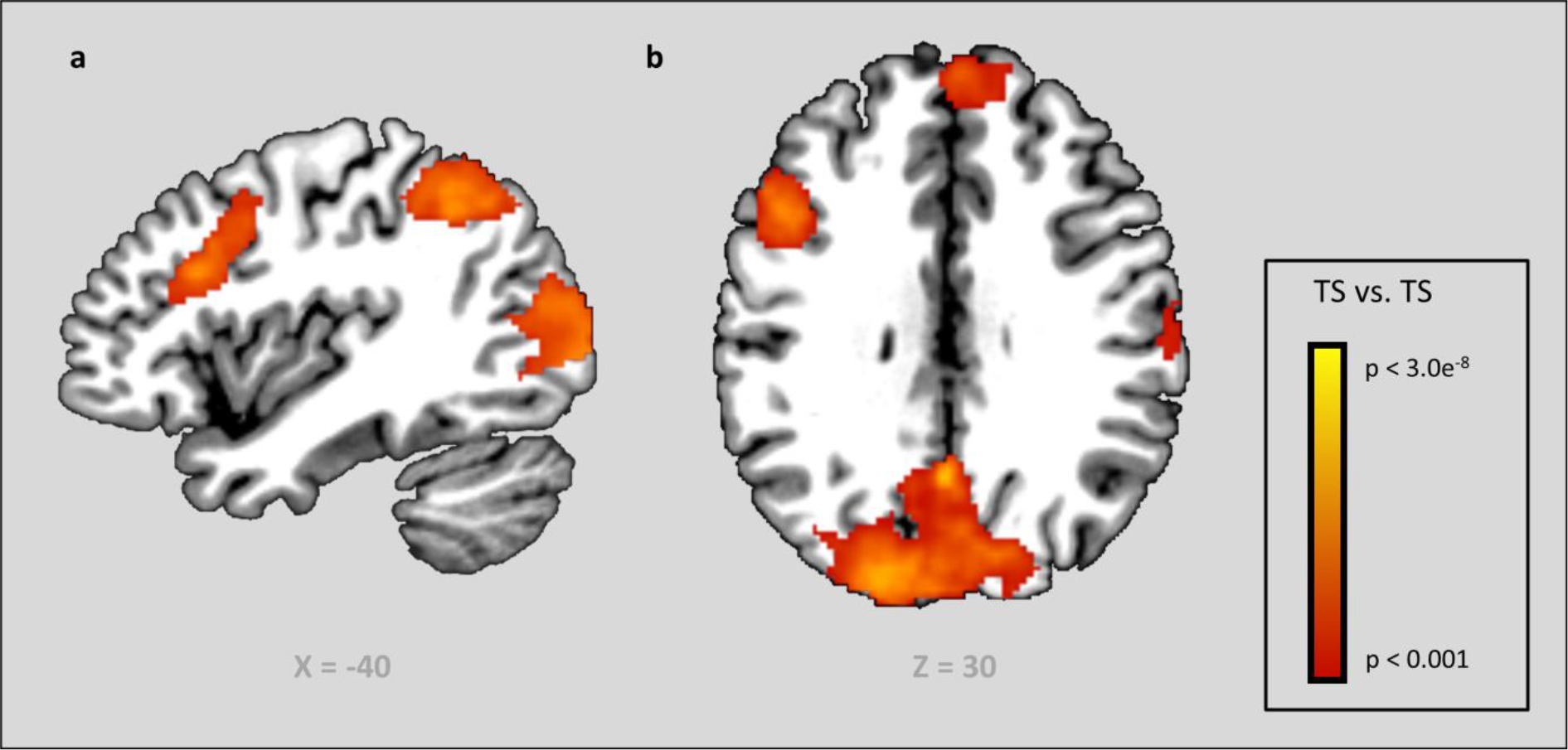
Searchlight results for classifying between the pairwise task sets. Clusters with significant (p<0.0001) above-chance classification accuracy for [task set vs. task set] can be found in bilateral occipital cortex, left lateral parietal cortex, right middle frontal gyrus, medial prefrontal cortex and left ventrolateral prefrontal cortex.

To assess whether the regions identified as encoding task-set information in our first-level analysis engaged in co-active or conjunctive representation, ROIs were defined based on the clusters described above. Due to a large interconnected cluster around parietal and visual cortex, the ROIs created for these regions were obtained by masking the cluster with an anatomical ROI derived from AAL labels (Tzourio-Mazoyer et al., 2002): left superior parietal and left inferior parietal labels were used for the parietal lobe mask, and bilateral middle occipital and calcarine sulcus labels for the visual cortex mask. Then, the most relevant features in these ROIs were identified (as described in the Methods section) for the 2nd-level classification. While our analyses consistently selected units that encoded task-set information in our synthetic data based on the strengths of network weights, feature selection based on weight strength may fail in some cases due to high noise or peculiarities in the structure of the data (Haufe et al., 2014). In particular, features required to maximally distinguish between two categories may in fact contain only noise, but nevertheless the optimal weight on that feature might exceed that of features that truly encode information. Therefore, in order to provide evidence that the features selected as the most relevant in our fMRI data do, in fact, encode task-related information, the first-level analysis for these ROIs was conducted again using only the features selected. Classification accuracy was similar when only the most-relevant features were used relative to classification done over the entire feature set (dlPFC mean = 59.37%, SE = 1.24%; mPFC mean = 54.52%, SE = 0.86%; parietal cortex mean = 59.24%, SE = 0.94%; visual cortex mean = 77.52%, SE = 1.50%). If the signal for the most-relevant features consisted purely of noise, classification accuracy using only the selected features would have been at chance. We can therefore rule out the possibility that the features identified as being most important to task-set classification were selected due to noise or the structure of the data.

Using the clusters defined and validated above, the 2nd-level classification accuracies for most- and least-important features were independently averaged across voxels within those ROIs for each subject (figure 9). A paired-samples t-test for each ROI indicated that, for lPFC, classification accuracy was greater for least-important voxels than most-important voxels, consistent with the co-activation account (p = 0.002, t(22) = −3.43). Parietal cortex also showed a trend towards co-activation, but the difference in most-important and least-important voxels failed to reach significance (p = 0.053, t(22) = −2.05). In contrast, classification accuracy was greater for most-important vs. least-important voxels in the occipital ROI, consistent with the conjunction account of representation (p < 0.001, t(22) = 6.14).

**Fig. 9.**
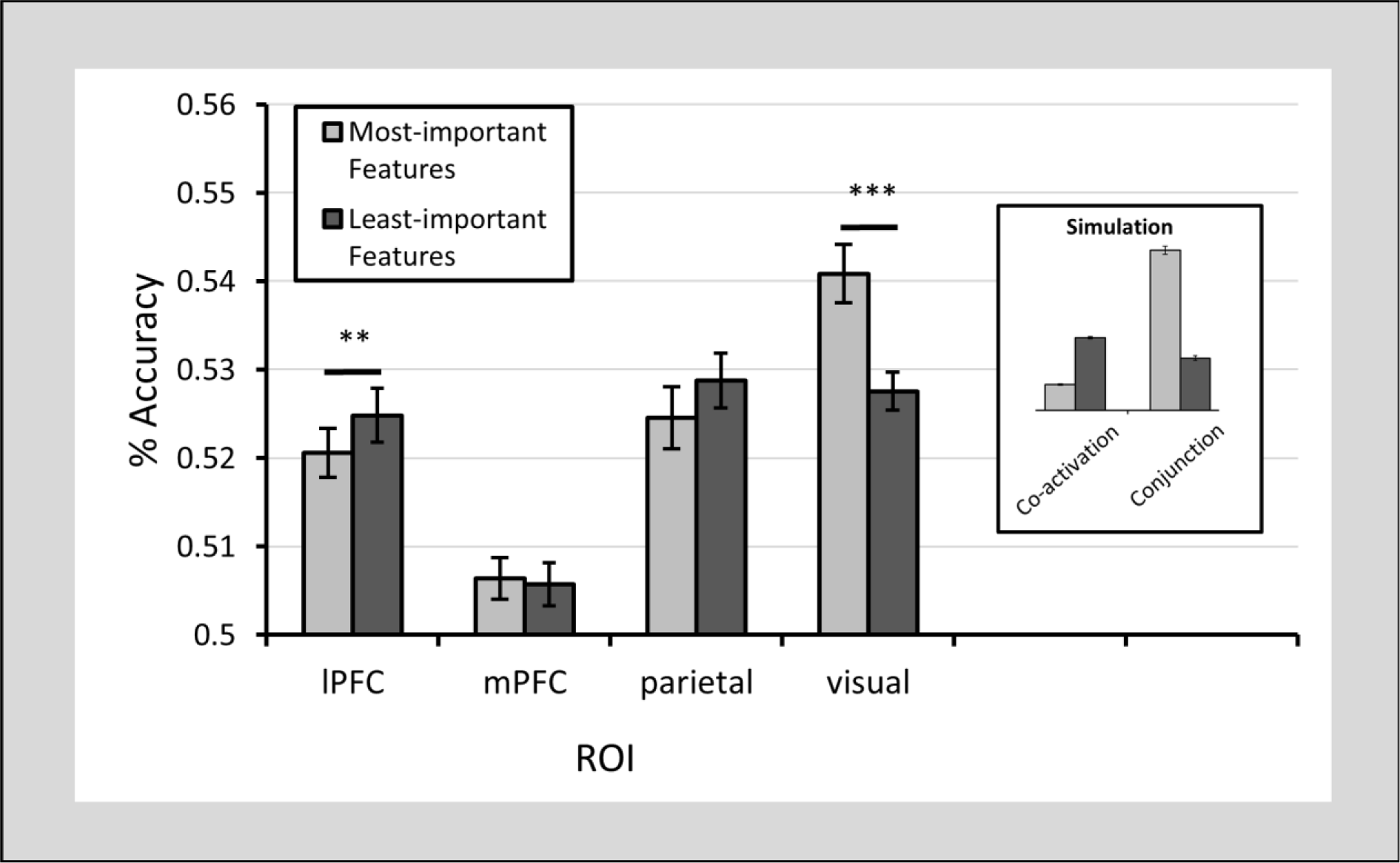
Results of the ROI analysis. Error bars show standard errors of the mean.

A full brain searchlight using the multi-level MVPA analysis (Figure 10 & Table 2) recapitulates our ROI analyses, showing classification in lPFC and parietal cortex consistent with co-activation, while bilateral visual cortex is consistent with conjunctive representation.

**Fig. 10.**
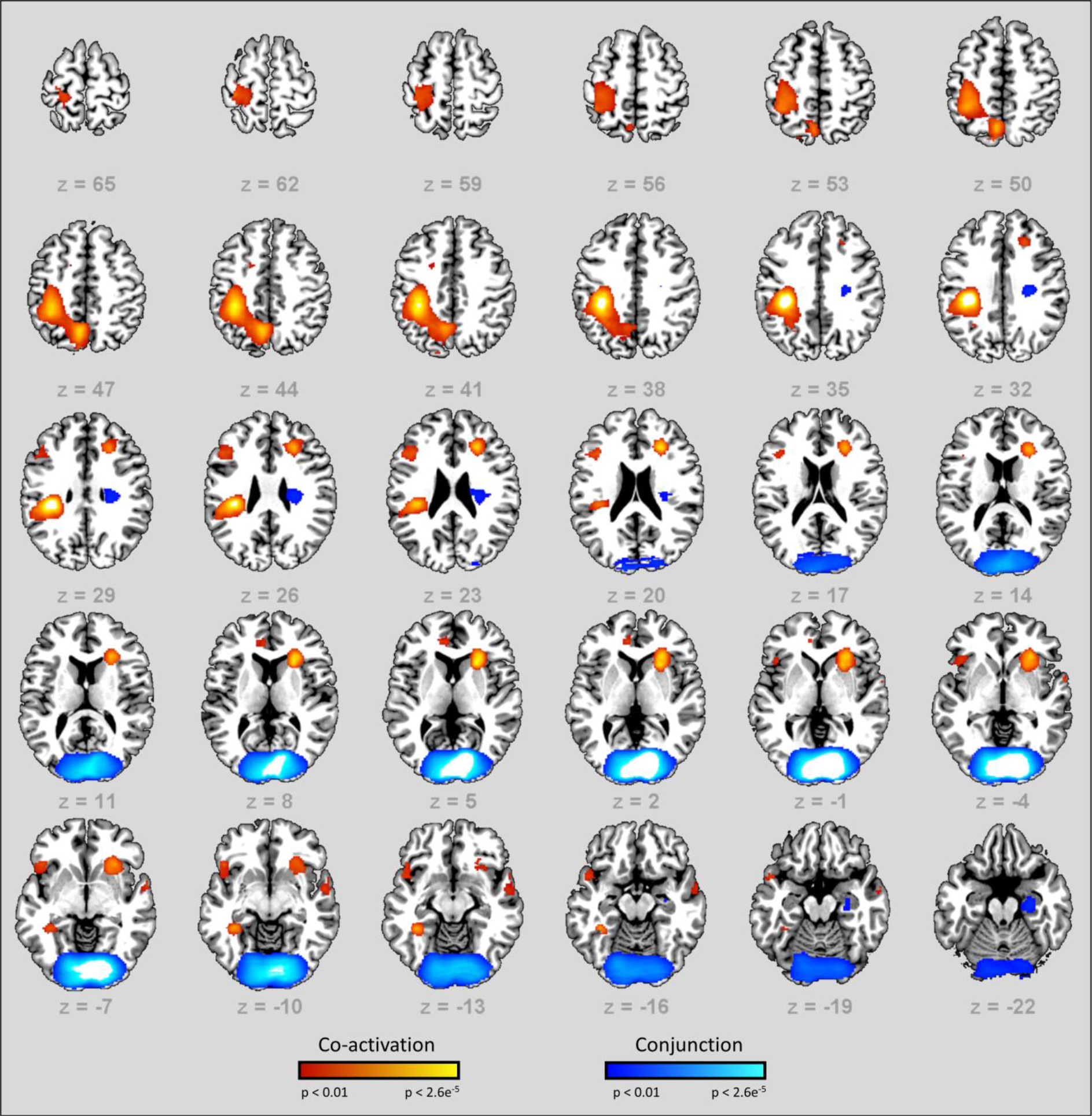
Results of the multi-level MVPA searchlight analysis. Blue represents the [most-important voxels > least-important voxels] contrast (*conjunction*), while orange is [least-important voxels > most-important voxels] (*co-activation*). The maps were thresholded at p < (uncorrected), with a voxel extent of 100.

**Table 2.** Summary of the decoding clusters for the multi-level MVPA searchlight analysis.

|  | Local Maxima | Cluster | Peak | Cluster-level |
| --- | --- | --- | --- | --- |
| Area | MNI | Size | T | p(uncorrected) |
| *Most-important Voxels > Least-important Voxels* |  |  |  |  |
| Calcarine sulcus | 14 -92 2 | 14826 | 11.13 | 0.000 |
| Right parahippocampal gyrus | 24 -16 -28 | 740 | 3.53 | 0.024 |
| Right precentral gyrus | 28 -20 38 | 720 | 3.43 | 0.026 |
| *Least-important Voxels > Most-important Voxels* |  |  |  |  |
| Left inferior parietal | -40 -30 40 | 8222 | 5.74 | 0.000 |
| Right anterior insula | 26 20 4 | 2657 | 4.00 | 0.000 |
| Left inferior orbitofrontal | -50 24 12 | 675 | 3.64 | 0.030 |
| Left inferior frontal triangularis | -42 26 20 | 576 | 3.40 | 0.043 |

## Discussion

In this study we used fMRI in combination with a novel multi-level MVPA approach to investigate “where” and “how” task sets are represented in the brain. Specifically, we sought to answer the question of whether areas involved in maintaining task-set representations did so in a co-active or conjoined manner. Our results, using classical univariate analysis as well as typical MVPA approaches, replicate standard findings from the WM literature. Furthermore, by incorporating feature extraction into our multi-level MVPA approach, as well as an experimental design explicitly formulated to take advantage of this novel method, we were able to identify specific regions in the brain involved in co-active and conjunctive representation of task sets.

### Univariate & Basic Searchlight results

In our univariate analyses, we observed elevated maintenance activity (task-sets vs. control trials) in several regions, including lPFC and visual cortex. Elevated lPFC activation is routinely implicated in studies on WM and has been shown to be involved in task set and load effects (Rottschy et al., 2012), and rule retention (Courtney et al., 1997; Clayton E. Curtis & D’Esposito, 2003; Smith & Jonides, 1999). Our observation of elevated maintenance-period activity for task sets in visual cortex is an equivocal finding. Some studies have reported no sustained levels of activity for visual cortex during retention, but do find differences in distributed patterns of activity corresponding to stimulus-specific features, which can be decoded with high accuracy (Harrison & Tong, 2009; Xing, Ledgeway, McGraw, & Schluppeck, 2013). Other studies do report increased sustained activity (Kastner, Pinsk, De Weerd, Desimone, & Ungerleider, 1999; Silver, Ress, & Heeger, 2007) and this has been attributed to maintained attention processes (Offen, Schluppeck, & Heeger, 2009).

We additionally observed increased maintenance-related activity for control trials vs. task set trials in left and right parietal cortex. Parietal cortex is often observed in studies of WM, and elevated activity in the region has been found to be correlated with active memory storage (C. E. Curtis, 2006; Dumontheil et al., 2010; Ester, Sprague, & Serences, 2015). Alternatively, since control trials require a response on the first stimulus after the cue, while for task set trials this occurs after the second stimulus after the cue, this elevated parietal cortex activity could also be related to task preparation processes (Phillips, Velanova, Wolk, & Wheeler, 2009; Rushworth, Johansen-Berg, Göbel, & Devlin, 2003).

### Standard MVPA analysis

To further investigate areas that may be involved in representing task set information, we turn to our multi-level MVPA approach. The first level of our approach is equivalent to typical MVPA analyses in which voxels within a specified anatomical region are used as input to a classifier. In this first level, a searchlight pairwise classification of task sets showed significant classification accuracy in visual cortex, as well as areas of the fronto-parietal WM network, including lPFC, mPFC and parietal cortex regions that are crucial to stimulus and rule encoding (Zhang, Kriegeskorte, Carlin, & Rowe, 2013).

### MVPA with feature selection

At the second level of our multi-level MVPA analysis, we selected voxels identified as important in our first level, pairwise classification, and used these as input to a second classifier which was trained on two novel, composite categories, one of which was composed of two task-sets that had been used to identify relevant voxels, and the other composed of the remaining task set and control trials. As outlined in our analysis of simulated data, this analysis predicts differential patterns of accuracy depending on the nature of the underlying representation. We observed classification accuracy in lPFC consistent with the co-activation hypothesis, i.e. classification accuracy was significantly lower when using the most-important voxels as input than when using the least-important voxels. This fits well with the finding that individuals can rapidly learn complex novel tasks through the transfer of practiced rule representations within lPFC to novel contexts. According to Cole et al. (2011), this transfer is made possible because rule representations in lPFC are compositional (i.e. co-active). That lPFC represents sets of rules co-actively is also consistent with previous work by Reverberi et al. (2012a) in which above-chance classification of task-sets in lPFC was achieved by training classifiers only on the simple rules they were composed of. While this shows that their compound rules follow a compositional structure in lPFC, the design described in Reverberi et al. (2012a) used compound rule trials interspersed with single rule trials that made up the compound rules, potentially influencing the participants to represent task sets as a composition of the single rules. In our study, however, rules were only presented as part of a larger task set, and single rules were never shown individually. However, it is possible that our task also favors co-active representations by requiring the counting of feature mismatches. Future experiments could use symbolic cues instead of the word cues used here to see if this biases the results. Furthermore, in contrast to Reverberi et al. (2012a), we additionally found tentative evidence for co-activation in a large region of lateral parietal cortex (Fig. 8, 9). While Reverberi et al. (2012a) did observe above-chance classification of task-sets in lateral parietal cortex; they were unable to decode these task sets from classifiers trained on their constituent rules. Given these discrepant findings, future work should more directly investigate the nature of representation in parietal cortex during WM.

While we observed classification accuracy consistent with the co-activation hypothesis in lPFC and parietal cortex, regions frequently associated with maintenance and implementation of task-sets composed of multiple rules, only classification accuracy in visual cortex was consistent with the conjunction hypothesis. The question of what is being represented here is difficult to answer. One possibility is that activity in visual cortex is primarily related to stimulus-specific features such as, in the present study, the shape of the cue. Many studies indeed report that visual cortex contains information regarding simple visual features held in WM (Harrison & Tong, 2009; Serences, Ester, Vogel, & Awh, 2009; Xing et al., 2013) and that prefrontal regions such as lPFC modulate visual activity during WM (Sreenivasan, Curtis, & D’Esposito, 2014). In this case, co-active representations of task-sets maintained in prefrontal regions such as lPFC could serve as top-down signals used to enhance selectivity of high-level conjunctive features in visual cortex. However, this does not necessarily entail that visual cortex is only doing conjunction. Based on our simulations (see Methods), a mixture of conjunctive and co-active representations would still show a conjunctive decoding pattern (Fig 7). Furthermore, it is unlikely that receptive fields in visual cortex are purely co-active or conjunctive (Liu et al., 2016). Future research should determine whether visual cortex does indeed exhibit both co-active and conjunctive representations.

Our finding that lPFC stores complex sets of rules in a co-active manner has implications for models of neurocognitive behavior, especially models that attempt to explain or simulate lPFC function. Oftefn, these models use independent, conjunctive coding for representations in lPFC (Collins & Frank, 2013; Holroyd & Yeung, 2012; Mack, Preston, & Love, 2017; O’Reilly & Frank, 2006; Rougier, Noelle, Braver, Cohen, & O’Reilly, 2005), while our results, and those of other studies (Reverberi et al., 2012a), are consistent with co-active encoding. Recent computational models of lPFC, such as the HER model (Alexander & Brown, 2015), suggest that task-set representations maintained in WM are co-active in nature, and specifically hypothesize distributed, co-active error representations within lPFC. While the results of present study are consistent with the HER model account, additional work is needed to more directly test whether the representations maintained in lPFC are specifically related to error representation. More generally, the evidence for co-active representation of task-sets provided here gives an intuitive explanation of how the brain is able to encode the enormous amount of rule combinations required for everyday behavior (Reverberi et al., 2012a).

## Conclusion

Using a multi-level MVPA approach in combination with fMRI, we found both “where” and “how” task sets are represented in the brain. Converging evidence from a variety of sources showed a distributed network of regions involved in representing task sets including lPFC, mPFC, parietal cortex and visual cortex. Further analyses showed the fronto-parietal regions of lPFC and possibly parietal cortex to be consistent with a co-active representational scheme, while visual cortex better fit the conjunction hypothesis. These findings support earlier work that shows compositional coding of task sets based on their constituent rules within lPFC (Reverberi et al., 2012a). The results shown here also illustrate the advantages of MVPA approaches, which allow us to answer questions beyond the scope of classical univariate analyses, and of feature selection, which has uses beyond dimensionality reduction and accuracy-boosting. Further research using feature selection in a similar manner as we did here in areas encoding co-actively could enable us to predict novel task set trials based on familiar task set trials composed from different combinations of rules. Additionally, the application of feature selection might be useful in elucidating the representational nature of other task variables such as error signals or object categories.

